# The SPARC complex defines RNAPII promoters in *Trypanosoma brucei*

**DOI:** 10.1101/2022.07.14.499694

**Authors:** Desislava P. Staneva, Stefan Bresson, Tatsiana Auchynnikava, Christos Spanos, Juri Rappsilber, A. Arockia Jeyaprakash, David Tollervey, Keith R. Matthews, Robin C. Allshire

**Affiliations:** Wellcome Centre for Cell Biology and Institute of Cell Biology, School of Biological Sciences, University of Edinburgh. Edinburgh EH9 3BF, Scotland, UK; Institute of Immunology and Infection Biology, School of Biological Sciences, University of Edinburgh, Edinburgh EH9 3JT, Scotland, UK; Institute of Biotechnology, Technische Universität, Gustav-Meyer-Allee 25, 13355 Berlin, Germany; The Francis Crick Institute, 1 Midland Road, London NW1 1AT, UK

**Author notes:** Co-corresponding authors: Keith Matthews, Robin Allshire.

## Abstract

Kinetoplastids are a highly divergent lineage of eukaryotes with unusual mechanisms for regulating gene expression. We previously surveyed 65 putative chromatin factors in the kinetoplastid *Trypanosoma brucei*. Our analyses revealed that the predicted histone methyltransferase SET27 and the Chromodomain protein CRD1 are tightly concentrated at RNAPII transcription start regions (TSRs). Here we report that SET27 and CRD1, together with four previously uncharacterized constituents, form the SET27 promoter-associated regulatory complex (SPARC), which is specifically enriched at TSRs. SET27 loss leads to aberrant RNAPII recruitment to promoter sites, accumulation of polyadenylated transcripts upstream of normal transcription start sites, and conversion of some normally unidirectional promoters to bidirectional promoters. Transcriptome analysis in the absence of SET27 revealed upregulated mRNA expression in the vicinity of SPARC peaks within the main body of chromosomes in addition to derepression of genes encoding variant surface glycoproteins (VSGs) located in subtelomeric regions. These analyses uncover a novel chromatin-associated complex required to establish accurate promoter position and directionality.

## INTRODUCTION

Kinetoplastids are a group of highly divergent eukaryotes that separated from the main lineage early in eukaryotic evolution. Reflecting this, kinetoplastids display many unusual molecular and biochemical features distinct from those of conventional eukaryotes. For example, gene expression regulation in these organisms is very different from conventional model eukaryotes such as yeasts, fruit flies, nematodes and vertebrates, all of which are members of the Ophistokont clade (Akiyoshi and Gull, 2013; Clayton, 2019; Keeling and Burki, 2019).

*Trypanosoma brucei* is a diploid kinetoplastid parasite transmitted to mammals by the tsetse fly in sub-Saharan Africa where it causes human sleeping sickness and animal nagana. In the mammalian host, bloodstream form (BF) trypanosomes have a surface coat composed of variant surface glycoprotein (VSG). Only one VSG is expressed at a time from a collection of ∼2000 distinct VSG genes and gene fragments, many of which are clustered near telomeres (Horn, 2014). *T. brucei* evades the mammalian immune system by occasionally switching to express a new VSG to which the host has not produced antibodies. Tsetse flies take up trypanosomes when they feed on infected mammals. In the fly midgut, *T. brucei* differentiates to the procyclic form (PF), a transition coupled to metabolic reprogramming and replacement of VSG with procyclins on the surface of these parasites (Matthews, 2005; Smith et al., 2017). Unusually, the active VSG and procyclin mRNAs are transcribed by RNAPI which is hypothesized to result from the high demand for these surface coat proteins at different stages of the *T. brucei* life cycle.

Most other *T. brucei* protein-coding genes are transcribed by RNAPII within long polycistrons from which individual mRNAs are processed using signals in adjacent 5′ and 3′ untranslated regions (UTRs; Benz et al., 2005; Kolev et al., 2010). Short spliced leader (SL) sequences are trans-spliced onto all pre-mRNAs, and this is mechanistically linked to the 3′ end processing of upstream genes (Matthews et al., 1994; Michaeli, 2011). Poly(A) tails are added to the 3′ end of all mRNAs, including VSG gene transcripts. Because of their polycistronic arrangement, RNAPII transcription of *T. brucei* genes is initiated at only ∼150 sites in a genome that encodes ∼9000 proteins (Berriman et al., 2005; Daniels et al., 2010). Such transcriptional start regions (TSRs) are analogous to the promoters of conventional eukaryotes and similarly exhibit bidirectional/divergent (dTSR) or unidirectional/single (sTSR) activity. TSRs are broad 5-10 kb regions marked by the presence of the histone variants H2A.Z and H2B.V and enriched for specific histone modifications, including methylation on H3K4 and H3K10, and acetylation on H3K23, H4A1, H4K2, H4K5 and H4K10 (Kraus et al., 2020; Siegel et al., 2009; Wright et al., 2010).

RNAPII transcription termination regions (TTRs) are marked by the presence of chromatin containing the H3.V and H4.V histone variants and the kinetoplastid-specific DNA modification base J (Schulz et al., 2016; Siegel et al., 2009). Approximately 60% of TTRs are located in regions where two neighbouring polycistrons end (convergent TTRs, cTTRs). The remaining 40% of TTRs are found between polycistrons oriented head-to-tail (single TTRs, sTTRs), and frequently coincide with RNAPI or RNAPIII transcribed genes.

Regulation of trypanosome gene expression is thought to occur predominantly post-transcriptionally via control of pre-mRNA processing, turnover and translation, suggesting a lack of promoter-mediated modulation. It was therefore surprising that a large number of putative chromatin regulatory factors exhibit enrichment in the vicinity of TSRs (Schulz et al., 2015; Siegel et al., 2009; Staneva et al., 2021). We previously identified two classes of TSR-associated factors. In chromatin immunoprecipitation assays, Class I factors, such as the predicted lysine methyltransferase SET27 and the Chromodomain protein CRD1, exhibit sharp peaks that coincide with the 5′ end of nascent pre-mRNAs. In contrast, Class II factors are distributed more broadly from the Class I peak and extend into the downstream transcription units (Staneva et al., 2021).

Here we set out to characterize the Class I factors SET27 and CRD1 which exhibit some of the most prominent peaks at RNAPII TSRs. Our previous affinity purifications and mass spectrometry identified four uncharacterized proteins that interact strongly with both SET27 and CRD1. We show that all six proteins associate with each other and that the four uncharacterized proteins also display sharp peaks at TSRs, coincident with sites where SET27 and CRD1 reside. Thus, these six proteins form the SET27 promoter-associated regulatory complex (SPARC). We further show that SPARC is required for accurate transcription initiation and/or promoter definition.

## RESULTS

### SET27 and CRD1 associate with proteins containing chromatin reader modules

We previously reported that YFP-tagged SET27 and CRD1 coimmunoprecipitate with each other and with JBP2, a thymidine hydroxylase involved in the synthesis of base J (DiPaolo et al., 2005; Kieft et al., 2007). In addition, both SET27 and CRD1 interacted with four uncharacterized proteins (Tb927.11.11840, Tb927.3.2350, Tb927.1.4250, Tb927.11.13820; Staneva et al., 2021). SET27 is predicted to contain a SET methyltransferase domain (Dillon et al., 2005) and a Zinc finger domain (Klug and Rhodes, 1987) whereas CRD1 has a putative methyl lysine binding Chromodomain (Paro, 1990; Singh et al., 1991; Supplemental Figure S1A). Among the four uncharacterized proteins, Tb927.11.11840 was named CSD1 because it contains a divergent Chromoshadow domain (Aasland and Stewart, 1995), normally found in association with Chromodomains. The uncharacterised protein Tb927.3.2350 has a predicted PHD finger histone methylation reader domain (Aasland et al., 1995), and was thus designated as PHD6. Tb927.1.4250 lacked strongly predicted domains but because it was enriched at promoters (see below), we named it Promoter Binding Protein 1 (PBP1). Finally, Tb927.11.13820 was designated PWWP1 due to its structural similarity to a PWWP domain of the human NSD2 histone methyltransferase, which binds methylated H3K36 (Arrowsmith and Schapira, 2019; Zhang et al., 2021). Overall, the presence of putative histone writer and reader domains suggests that these proteins may cooperate to regulate chromatin structure and gene expression.

### Identification of the TSR-associated SPARC complex in bloodstream form *T. brucei*

The enrichment of CSD1, PHD6, PBP1, PWWP1 and JBP2 with affinity-selected YFP-SET27 and YFP-CRD1 (Staneva et al., 2021) initially suggested that these seven proteins might form a single complex. To investigate this further, we YFP-tagged the four uncharacterized proteins and JBP2 at their endogenous gene loci in bloodstream form (BF) Lister 427 *T. brucei* cells. Immunolocalization was performed to determine the subcellular distribution of the putative SET27-CRD1 complex components. Strong nuclear signals were obtained for CRD1, CSD1, PHD6, PBP1 and PWWP1 whereas SET27 and JBP2 were found in both the nucleus and the cytoplasm of BF cells (Supplemental Figure S1B). We subsequently affinity purified each protein and identified its interacting partners via the same LC-MS/MS proteomics pipeline we previously used for YFP-SET27 and YFP-CRD1 (Staneva et al., 2021).This analysis showed that each protein displayed strong association with the other six (Figure 1A; Supplemental Table S1).

**Figure 1.**
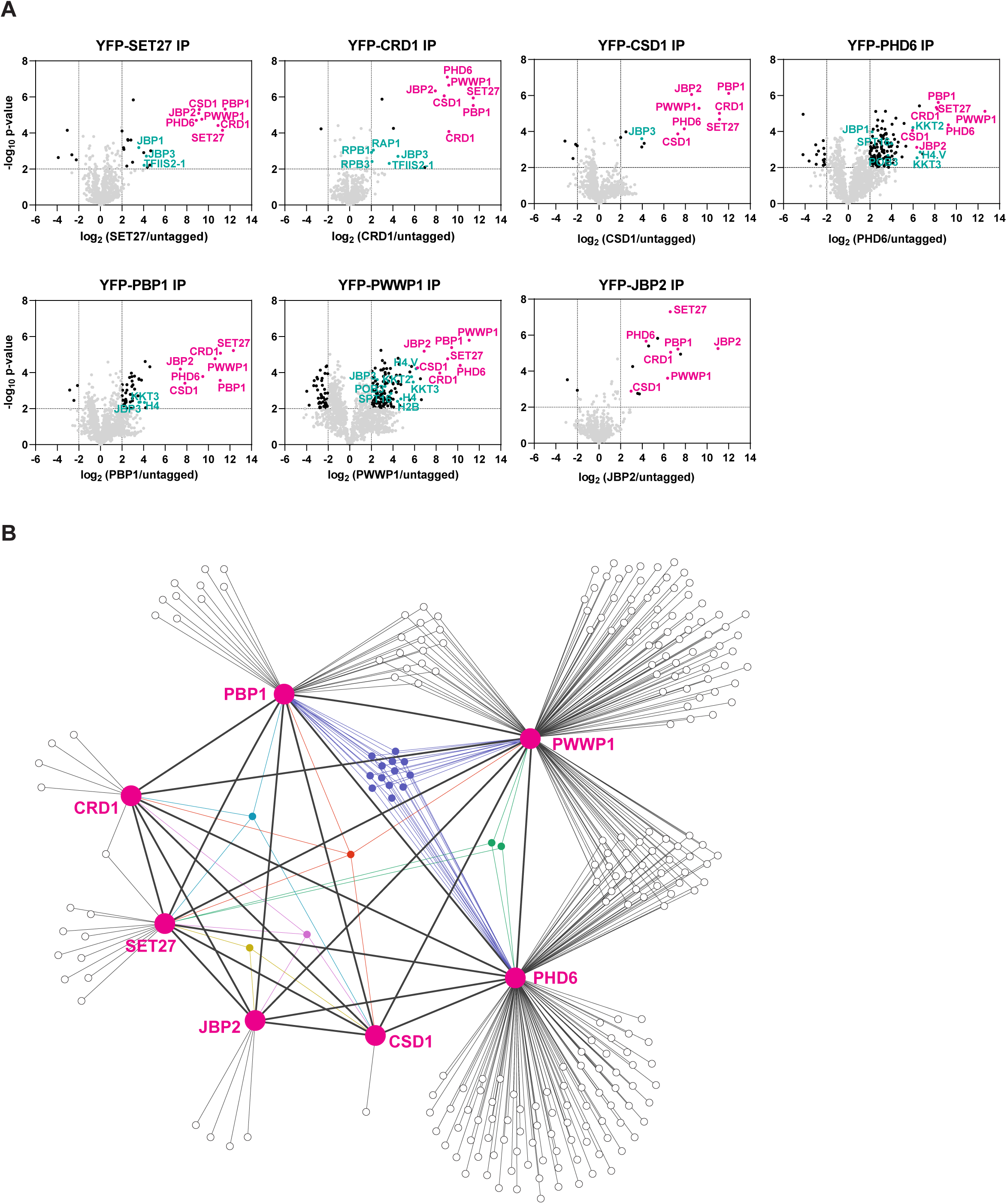
Identification of SPARC in bloodstream form *T. brucei*. **(A)** Proteins previously shown to be enriched in CRD1 and SET27 coimmunoprecipitations (co-IPs) were YPF-tagged, and analyzed by affinity selection and LC-MS/MS to identify protein interaction partners. The CRD1 and SET27 co-IPs are reproduced from Staneva et al., 2021. Volcano plots are based on 3 biological replicates for each sample. Cut-offs used for significance: log_2_ (tagged/untagged) >2 or <-2 and Student’s t-test p-value < 0.01. See Supplemental Table S1 for a complete list of proteins in each affinity selection. Putative SPARC complex subunits are shown in pink, and other proteins of interest are shown in teal. **(B)** Interaction network of the proteins enriched in the co-IP experiments shown in (A). SPARC components are connected with thick lines, while all other interactions are shown with thin lines. Proteins which interact with three or more SPARC components are represented in different colours. See Supplemental Table S2 for a complete list of shared and unique interactors in these co-IPs.

This group of seven represented the most highly enriched proteins in the affinity selections of SET27, CRD1, CSD1 and PBP1, being on average 8-fold more prevalent than any other proteins detected. However, several bait proteins recovered overlapping sets of additional proteins (Figure 1B; Supplemental Table S2). SET27, CRD1, CSD1, PBP1 and PWWP1 associated with JBP3, a base J binding protein involved in RNAPII transcription termination in *T. brucei* and *Leishmania* (Jensen et al., 2021; Kieft et al., 2020). PHD6 and PWWP1 affinity purifications were enriched for various histones (H2A, H2B, H3.V, H4, etc.) and kinetochore proteins (KKT2, KKT4, KKT8, etc.), and PHD6, PBP1 and PWWP1 each associated with the histone variant H4.V and the kinetochore protein KKT3. Recovery of histone proteins is consistent with the presence of putative histone-binding modules in CRD1, PHD6 and PWWP1. Additionally, CRD1, PHD6 and PWWP1 interacted with various RNAPII subunits, and both PHD6 and PWWP1 associated with the POB3 and SPT16 components of the FACT complex. FACT is involved in transcription elongation in eukaryotes (Belotserkovskaya et al., 2003), and in *T. brucei* it has also been linked to VSG repression (Denninger and Rudenko, 2014). Overall, our proteomic analyses indicate that SET27, CRD1, CSD1, PHD6, PBP1, PWWP1 and JBP2 are tightly associated, potentially forming a protein complex, with some components showing interactions with a wider group of proteins involved in transcriptional regulation (Figure 1B).

Previously we demonstrated that both SET27 and CRD1 are specifically enriched across a narrow segment of RNAPII TSRs in *T. brucei* BF cells (Staneva et al., 2021). We therefore performed ChIP-seq for the five other SET27/CRD1-associated proteins in BF parasites. CSD1, PHD6, PBP1 and PWWP1 each showed sharp peaks at TSRs, coincident with SET27 and CRD1 (Figure 2A and B). In contrast, we did not detect specific association of JBP2 with any genomic region. Thus, JBP2 is either not chromatin-associated, only transiently associates with chromatin or the fixation conditions used might be insufficient to crosslink it to chromatin. Consequently, despite the clear and robust affinity purification of JBP2 with the other six proteins in BF cells, we are unable to reliably designate JBP2 as a core component of the complex. Because SET27, CRD1, CSD1, PHD6, PBP1 and PWWP1 clearly associate with each other and are co-enriched over RNAPII promoter regions, we termed this six member complex the SET27-promoter-associated regulatory complex (SPARC).

**Figure 2.**
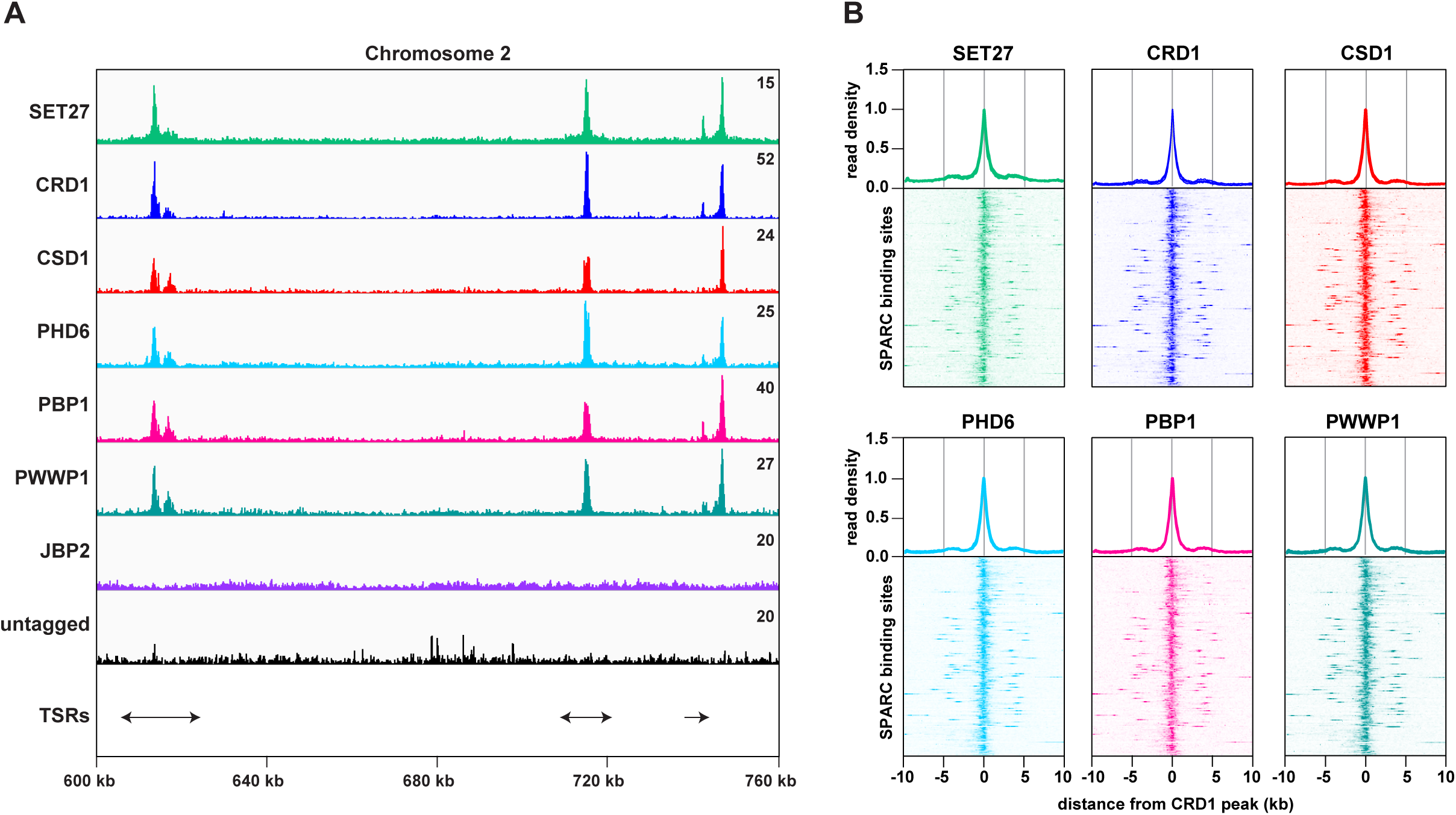
SPARC components target the same genomic loci in bloodstream form *T. brucei*. **(A)** A region of Chromosome 2 is shown with ChIP-seq reads mapped for the indicated proteins. A single replicate is shown for each protein. ChIP-seq data for CRD1, SET27 and the untagged parental cell line are reproduced from Staneva et al., 2021. ChIP-seq performed in wild type cells lacking any tagged protein (untagged) was included as a negative control. Tracks are scaled separately as fragments per million (the exact value is indicated in the top-right corner of each track). The position of single and double transcription start regions (sTSRs and dTSRs) are shown below with arrows indicating the direction of transcription. **(B)** Enrichment profiles for SPARC components. CRD1 is used as a reference because it has the most prominent peaks at TSRs. The metagene plots (*top*) show normalized read density around all CRD1 peak summits, with each replicate plotted separately. The heatmaps (*bottom*) show protein density around 177 individual CRD1 peaks. Each heatmap shows the average of at least two replicates.

### Procyclic form *T. brucei* cells also contain SPARC

To determine if SPARC exists with a similar composition in the distinct insect stage of the *T. brucei* life cycle, we YFP-tagged SET27, CRD1 and JBP2 at their endogenous loci in procyclic form (PF) parasites. In PF cells, JBP2 was predominantly cytoplasmic, whereas SET27 was strongly enriched in the nucleus. This contrasts with BF cells, where SET27 and JBP2 were detected in both the nucleus and cytoplasm (Supplemental Figure S1B). CRD1 localized to the nucleus in both developmental forms of *T. brucei*.

Affinity selections of YFP-tagged SET27 and CRD1 from PF cells showed that both proteins strongly associate with each other and with the other four core SPARC components. Indeed, CSD1, PHD6, PBP1 and PWWP1 were at least 6-fold more enriched in YFP-SET27 and YFP-CRD1 purifications relative to any other protein (Supplemental Figure S2A; Supplemental Table S1). However, in contrast to BF cells, affinity selected YFP-SET27 displayed only a weak association with JBP2 (50-fold less than other SPARC components) and YFP-CRD1 showed no interaction with JBP2 in PF trypanosomes (Supplemental Figure S2A; Supplemental Table S1). Similarly, core SPARC components were only weakly enriched with affinity selected YFP-JBP2 (∼2-fold less compared to other proteins). This weak association of SPARC subunits with JBP2 in PF cells is consistent with its distinct cytoplasmic localisation relative to core components of the complex (Supplemental Figure S1B).

We next performed ChIP-seq to map the genome-wide binding sites of YFP-tagged SET27, CRD1 and JBP2 in PF cells. As in BF cells, SET27 and CRD1 were both localized to RNAPII TSRs, whereas JBP2 was not chromatin-associated (Supplemental Figure S2B).

These analyses in PF cells indicate that six core components, SET27, CRD1, CSD1, PHD6, PBP1 and PWWP1, form the main SPARC complex in both BF and PF cells. The ancillary factor JBP2 was largely absent from PF cell nuclei and only very weakly associated with other SPARC components in PF cells, underscoring its distinct behaviour.

### Loss of SET27 disrupts SPARC formation

The strong association of SPARC with RNAPII TSRs in both BF and PF *T. brucei* cells suggested that it might have a role in regulating transcription. To test this, we attempted to delete the genes encoding SPARC components in BF cells. We successfully generated cell lines lacking the core subunit SET27 (*set27*Δ/Δ) and the auxiliary subunit JBP2 (*jbp2*Δ/Δ). However, we were unable to obtain cell lines completely null for any other SPARC component, suggesting these may be essential for viability. To confirm the *SET27* and *JBP2* gene deletions, we generated RNA-seq libraries using poly(A)-selected RNA. Cells lacking either one (Δ/+) or both alleles (Δ/Δ) showed a partial and complete loss of *SET27* and *JBP2* mRNA, respectively (Figure 3A; Supplemental Figure S4A). Importantly, transcripts derived from neighbouring genes were unaffected, demonstrating that deletion of the *SET27* or *JBP2* genes did not alter expression of other genes in their respective polycistrons.

**Figure 3.**
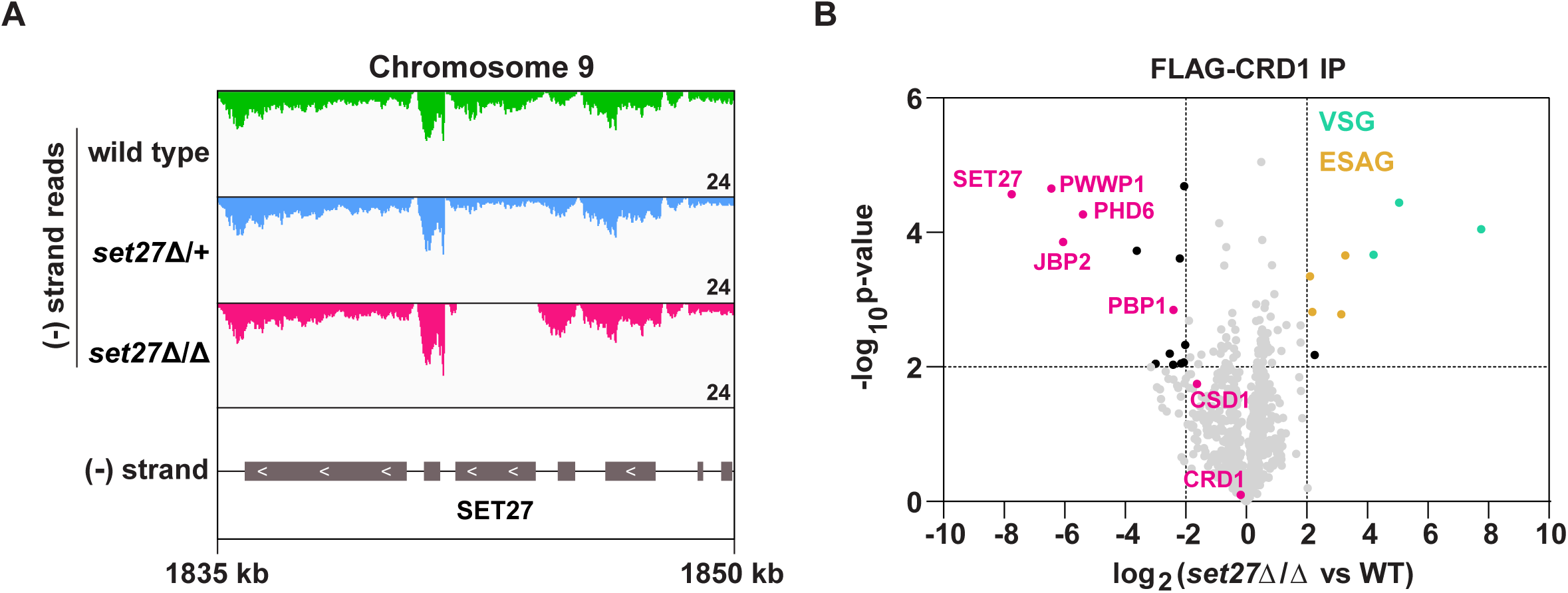
SPARC integrity is compromised in bloodstream form *T. brucei* lacking SET27. **(A)** Tracks showing the distribution of RNA-seq reads in the vicinity of the *SET27* gene in wild type, *set27*Δ/+ and *set27*Δ/Δ cells. All tracks are scaled identically (number of reads shown in the bottom right corner of each track). **(B)** Affinity selection of FLAG-CRD1 from wild type and *set27*Δ/Δ cells. The volcano plot is based on 3 biological replicates for each sample. Cut-offs used for significance: log_2_ (WT vs *set27*Δ/Δ) >2 or <-2 and Student’s t-test p-value < 0.01. See Supplemental Table S1 for a complete list of proteins in this affinity selection. SPARC components are shown in pink, VSGs in green and ESAGs in yellow.

Initial phenotypic characterization of cells lacking SET27 or JBP2 showed that *set27*Δ/Δ cells grew substantially slower than wild type or *set27*Δ/+ cells (Supplemental Figure S3A). In contrast, *jbp2*Δ/Δ cells grew slightly faster than wild type cells (Supplemental Figure S4B). The basis for these differences in growth rates remains to be determined.

To test if SET27 is required for the integrity of SPARC, we FLAG-tagged CRD1 at its endogenous locus in wild type and *set27*Δ/Δ cells. Next, we performed affinity selections of FLAG-CRD1 from both cell lines followed by mass spectrometry to detect potential differences in interacting proteins. In the absence of SET27, FLAG-CRD1 showed greatly reduced association with all other SPARC components (Figure 3B; Supplemental Table S1). Thus, we conclude that SET27 is pivotal for SPARC assembly or stability, as other subunits dissociate in its absence.

### SET27 is required for correct transcription initiation in bloodstream form *T. brucei*

Because SPARC is enriched over transcription start regions, we tested whether SET27 is required for accurate transcription initiation. In *T. brucei*, transcription initiates from either unidirectional or bidirectional promoters. All regions annotated as sTSRs in the trypanosome genome correspond to unidirectional promoters which are associated with a single SPARC binding site. Regions annotated as dTSRs exhibit either one or two SPARC peaks depending on whether the promoter is bidirectional or there are two nearby unidirectional promoters from which transcription initiates in opposite directions (for examples, see Figures 4 and 5). In wild type and *set27*Δ/+ cells, transcription typically initiated coincident with, or just downstream from, the peak of SPARC binding (Figure 4A-C). Strikingly, this pattern was lost in *set27*Δ/Δ cells. At all unidirectional promoters, the transcription start site shifted ∼1-5 kb upstream of its normal position in wild type cells (Figure 4A, *left*), though the extent of this defect varied between promoters in terms of both distance from the wild type start site and RNA amount. Additionally, at 14 out of 54 normally unidirectional promoters, transcription initiated in both directions in *set27*Δ/Δ cells, effectively converting them to bidirectional promoters (Figure 4A, *right*). These altered transcription patterns were also clearly evident in a metagene analysis of unidirectional promoters (Figure 4C). Data from dTSRs was more difficult to interpret because in these regions one cannot distinguish between an upstream and an antisense transcriptional defect. Nonetheless, there was strong evidence that SET27 also contributes to accurate transcription initiation from dTSRs (Figure 4B and C). To determine if the transcriptional phenotype we observed was specifically due to SET27 loss, we restored the wild type *SET27* gene to its endogenous locus by homologous recombination (Supplemental Figure S3B). We obtained two rescue clones which grew slower than wild type but faster than *set27*Δ/Δ cells (Supplemental Figure S3A). Importantly, the elevated level of transcription within two TSRs observed in *set27*Δ/Δ cells was completely reversed when the *SET27* gene was restored (Figure 4D).

**Figure 4.**
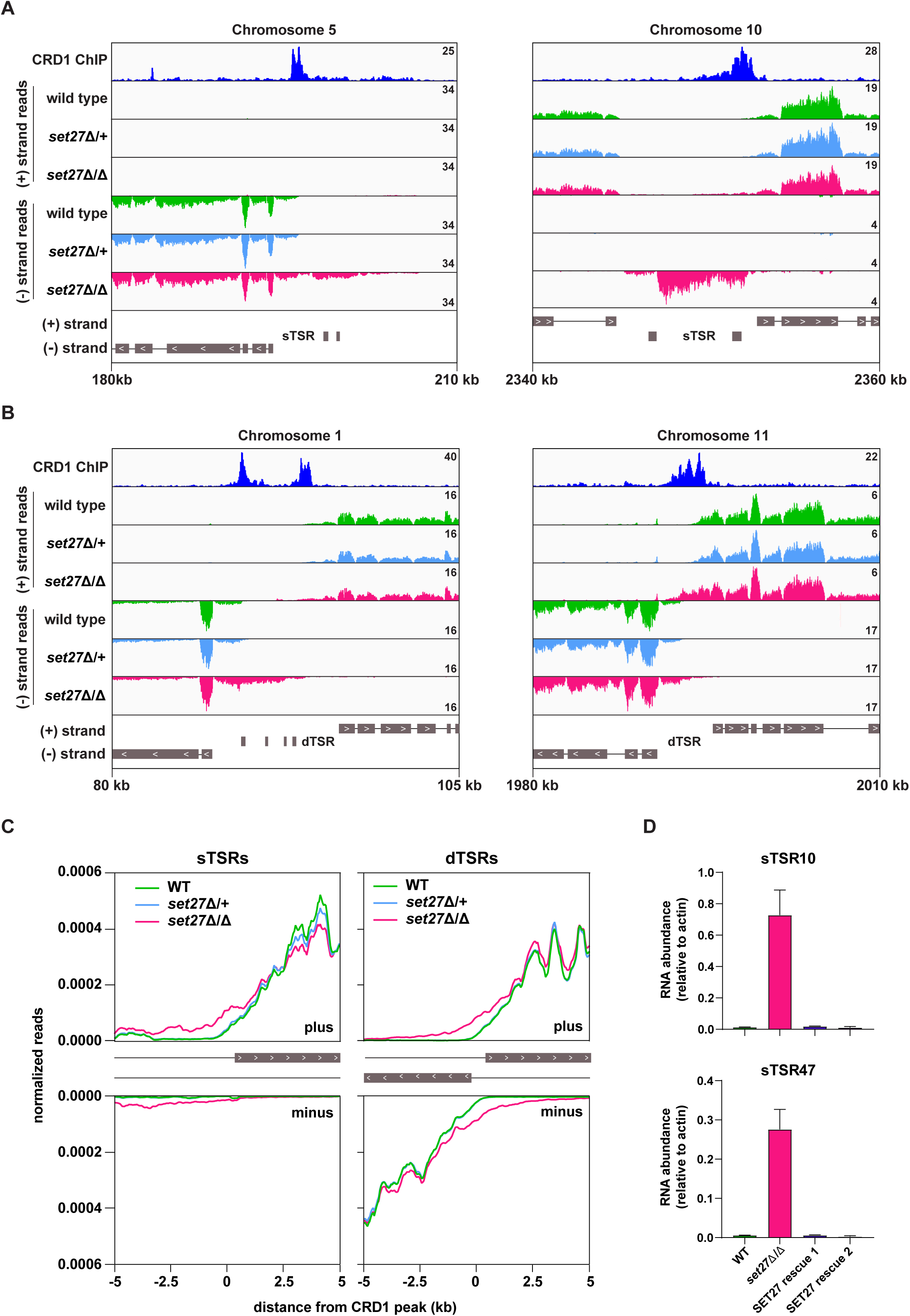
Transcription initiation is dysregulated in the absence of SET27. **(A)** Tracks showing the distribution of RNA-seq reads in the presence (wild type and set27Δ/+) or absence of SET27 (*set27*Δ/Δ) around selected unidirectional sTSR promoters. CRD1 ChIP (*top track*) is included to mark the position of SPARC sites. ORFs are indicated by grey boxes, and directionality is shown with inset white arrows. Genes present within a single polycistron are connected with a thin black line. Hypothetical protein-coding genes annotated within each TSR region are not connected to neighboring polycistrons. **(B)** As in (A), but for dTSR promoters. **(C)** *Left*: metaplots showing the distribution of RNA-seq reads around 33 SPARC sites marking unidirectional sTSR promoters. For clarity, we excluded SPARC sites present within 5 kb of a different SPARC site. Transcription in the forward direction is shown in the upper panel, and transcription in the reverse direction is shown below. *Right*: metaplots showing the distribution of RNA-seq reads surrounding all 120 SPARC sites present at dTSR promoters. **(D)** RT-qPCR of RNA originating from sTSR10 and sTSR47 in wild type, *set27*Δ/Δ and two SET27 rescue clones. Error bars: Standard Deviation (SD) of 2 biological replicates.

**Figure 5.**
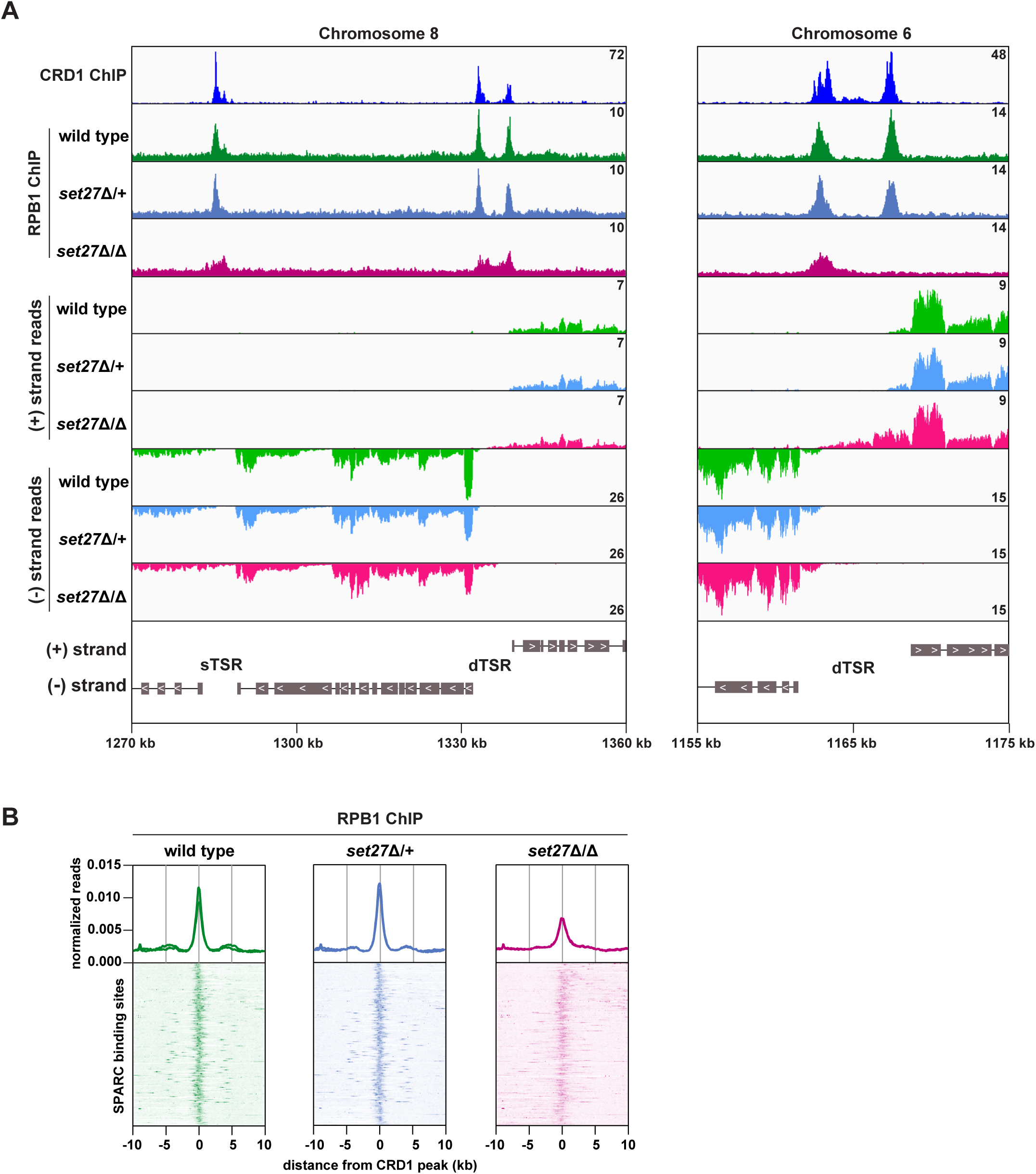
SET27 is required for full RNAPII recruitment to transcription start regions. **(A)** Tracks showing the distribution of YFP-tagged RPB1, the largest subunit of *T. brucei* RNAPII, across selected genomic windows following ChIP-seq. CRD1 ChIP (*top*) and RNA-seq (*bottom*) tracks are included for comparison. **(B)** RPB1 enrichment profiles. CRD1 is used as a reference because it has the most prominent peaks at TSRs. The metagene plots (*top*) show normalized read density around all SPARC sites, with each replicate plotted separately. The heatmaps (*bottom*) show RPB1 density around 177 individual SPARC sites. Each heatmap shows the average of at least 2 replicates.

In contrast to SET27, loss of JBP2 did not result in any distinct changes to the transcriptional profile around TSRs (Supplemental Figure S4C). This is consistent with our ChIP-seq data showing that JBP2 does not selectively associate with promoter regions (Figure 2), and again indicates that its function is distinct from core SPARC components. Collectively, these results suggest that SPARC contributes to the accurate positioning of transcription start sites and to normal transcription directionality.

### The distribution of RNAPII is altered in cells lacking SET27

If promoter positioning is indeed affected by SET27 loss, then RNAPII occupancy within TSRs would also be expected to change. To directly test this possibility, we performed ChIP-seq for YFP-RPB1 (the largest RNAPII subunit) in wild type, *set27*Δ/+ and *set27*Δ/Δ BF *T. brucei* cells. In wild type and *set27*Δ/+ cells, RPB1/RNAPII enrichment was clearly coincident with SPARC peaks (represented by CRD1; Figure 5). Notably, this pattern was significantly altered in *set27*Δ/∆ cells, with RPB1/RNAPII peaks becoming less defined and, in some instances, completely lost (Figure 5A). Comparison with RNA-seq data showed that the broader RPB1/RNAPII signal in *set27*Δ/Δ cells coincides with transcript accumulation upstream of the major transcription initiation site in wild type cells (Figure 5A). Metagene analysis confirmed that the RPB1/RNAPII peaks are generally reduced and broader in *set27*Δ/Δ cells relative to *set27*Δ/+ or wild type cells (Figure 5B). Thus, we conclude that SET27 is required for accurate RNAPII transcription initiation.

### SET27 represses the expression of VSG and transposon-derived genes

The finding that transcription initiation and RNAPII occupancy is altered in *set27*Δ/Δ cells prompted us to determine if these changes affect the expression of any mRNAs. Principal component analysis (PCA) was employed to compare the global transcriptional landscapes of wild type, *set27*Δ/+ and *set27*Δ/Δ cells (Figure 6A). Wild type and *set27*Δ/+ replicates clustered together while *set27*Δ/Δ replicates were clearly distinct, indicating that gene expression is aberrant in cells lacking SET27. Analysis of differentially expressed genes (DEGs) in wild type versus *set27*Δ/Δ cells revealed widespread transcriptional induction of otherwise silent or weakly-expressed genes (Figure 6B and C). Almost 100 mRNAs were significantly upregulated at least two-fold (Figure 6B; Supplemental Table S3). Conversely, only the *SET27* mRNA was reduced, due to its gene deletion (Figure 6B; Supplemental Table S3). Gene ontology analysis of the upregulated gene set revealed strong enrichment for normally silent VSG genes (Figure 6B-D). Additionally, we observed derepression of SLACS retrotransposons which reside within the spliced leader gene cluster, as well as derepression of reverse transcriptase and RNase H genes derived from the more widely distributed ingi retrotransposons (Figure 6B-D). Expression site-associated genes (ESAGs; transcribed together with the active VSG) were also upregulated in *set27*Δ/Δ cells (Figure 6B and C). However, the ESAG gene category was not significantly enriched overall among the DEGs (Figure 6D).

**Figure 6.**
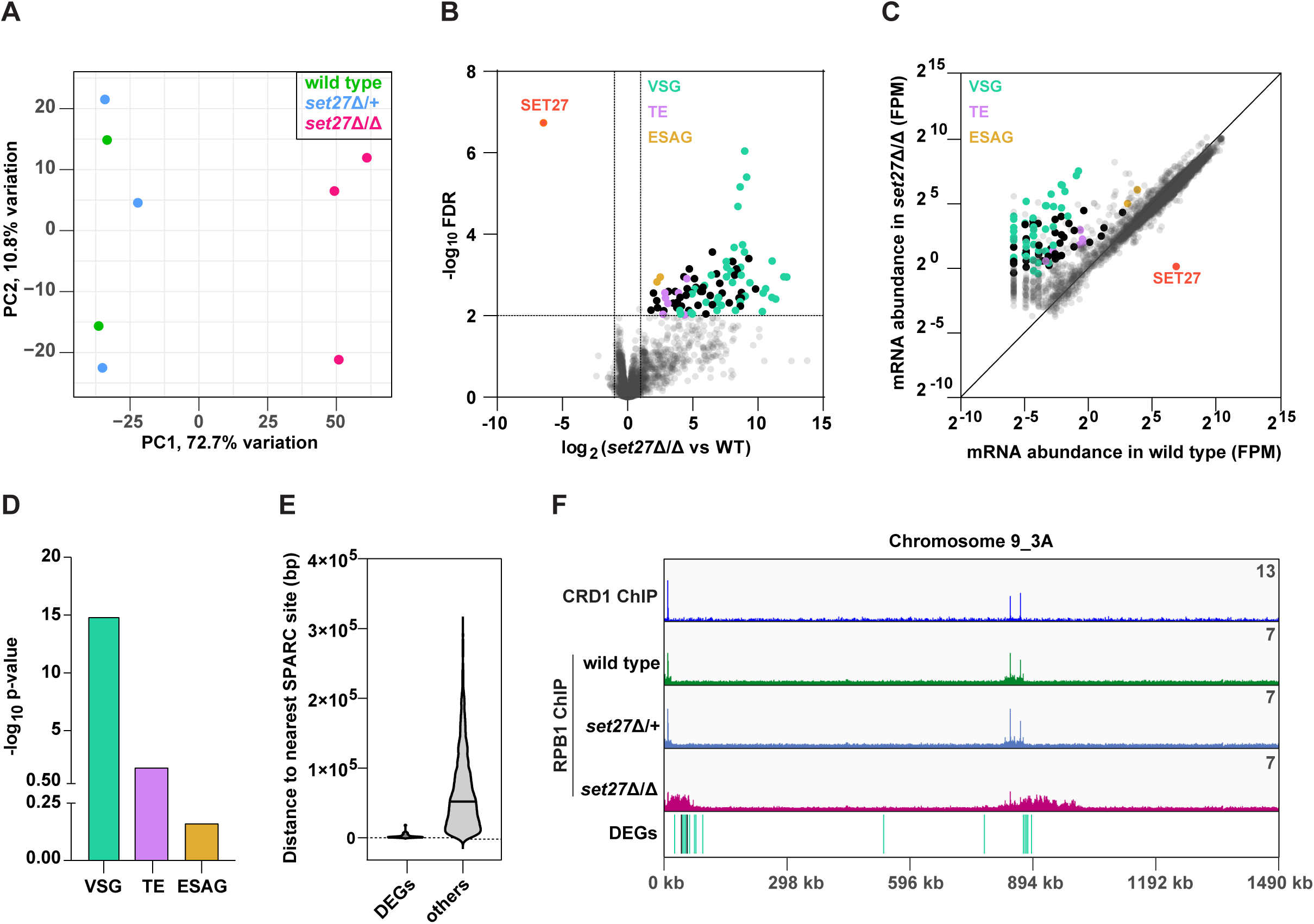
SET27 deletion derepresses VSGs and transposon-associated genes. **(A)** PCA comparing mRNA expression (n=8540) in wild type, *set27*Δ/+, and *set27*Δ/Δ cells. **(B)** Volcano plot showing differentially expressed genes between wild type and *set27*Δ/Δ cells. Cutoffs used for significance: log_2_ (tagged/untagged) >1 or <-1 and FDR <0.01. VSGs are coloured in green; SLACS retrotransposable elements and ingi-derived reverse transcriptase and RNase H genes are in lavender; ESAG genes are in yellow; other DEGs are in black; and non-DEGs are in gray. See Supplemental Table S3 for a complete list of DEGs. **(C)** Scatter plot of mRNA abundance normalized as FPM (fragments per million) in wild type versus *set27*Δ/Δ cells. The diagonal marks the position of genes with equal expression in the wild type and *set27*Δ/Δ cell lines. The colour scheme is the same as in (B). **(D)** Gene ontology enrichment among upregulated mRNAs. P-values were calculated using Fisher’s exact test. **(E)** Violin plot showing the distance between the 5**′** end of each differentially expressed gene (DEG) and the nearest SPARC site. Non-DEGs (‘other’) are included as a comparison. Only genes found in the main body of chromosomes were included in this analysis. Subtelomeric genes were excluded because they frequently recombine, and their precise location is uncertain. **(F)** The Chromosome 9_3A subtelomeric contig showing ChIP-seq reads for CRD1 and RPB1. The locations of DEGs are shown below. The colour scheme is the same as in (B).

To determine if the DEGs identified as being upregulated in *set27*Δ/Δ cells share any particular feature that might explain their sensitivity to SPARC loss, we mapped their genomic location relative to the nearest SPARC peak. Genes whose exact genomic location varies, such as VSG genes that are located at subtelomeric regions and can frequently recombine to new locations, were excluded from this analysis. Notably, upregulated genes typically resided within 2.5 kb of a SPARC peak, compared to 65 kb for the average, unaltered gene (Figure 6E).

A sizable subset of DEGs were VSG genes located at the ends of chromosomes adjacent to telomeres. Interestingly, upon manual inspection, we noticed that a large number of these induced subtelomeric VSG transcripts (16 out of 37) were derived from two locations on chromosome 9 near the right telomere. In both cases these VSG gene transcripts arise downstream of SPARC peaks but are not immediately adjacent to them (Figure 6F). In the absence of SET27, RPB1/RNAPII displayed increased chromatin association over a much larger region downstream of each SPARC peak: ∼60 kb and ∼130 kb regions containing 11 and 5 differentially expressed VSGs, respectively (Figure 6F). We also observed upregulation of two non-VSG genes, indicating that general derepression occurs across this subtelomeric region.

These analyses demonstrate that SPARC is required to restrict the expression level of genes in the immediate vicinity of TSRs located within the main body of *T. brucei* chromosomes as well as genes, particularly VSGs, located further downstream of SPARC sites within subtelomeric regions.

## DISCUSSION

In this study, we identified SPARC, a promoter-associated protein complex in both bloodstream and procyclic forms of *T. brucei*, comprising SET27, CRD1, CSD1, PHD6, PBP1 and PWWP1. Several SPARC components showed homology to known histone mark reader and writer domains, including SET, Chromo, Chromoshadow, PHD and PWWP, indicating that the association of SPARC with chromatin is mediated through binding to specific histone modification(s). JBP2 also copurified with SPARC components but it appears to be functionally distinct with respect to its chromatin binding and impact on transcription initiation. In contrast to a previous report (DiPaolo et al., 2005), we detected YFP-JBP2 and identified untagged JBP2 as a YFP-SET27 interactor in PF cells, indicating that JBP2 is expressed in procyclic form *T. brucei*.

A chromatin-associated complex containing homologs of SET27, CRD1, CSD1 and PBP1 was recently identified in *Leishmania* (Jensen et al., 2021), showing that SPARC-related complexes are also present in other kinetoplastids, albeit with potentially altered composition. The *Leishmania* complex subunits interacted strongly with JBP3, a DNA base J binding protein, and deletions of JBP3 were shown to result in transcriptional read-through downstream of RNAPII termination sites in both *T. brucei* and *Leishmania* (Jensen et al., 2021; Kieft et al., 2020). Similarly, we detected JBP3 in affinity purifications of most core SPARC components. However, neither SET27 nor JBP2 deletions resulted in transcription termination defects, indicating that JBP3 function is distinct from SPARC.

While most SPARC subunits appear to be essential for viability, we were able to delete both alleles of the *SET27* gene, leading to dissociation of the other complex subunits. In the absence of SET27, polyadenylated transcripts in the vicinity of SPARC binding sites were upregulated as a result of transcript accumulation upstream of the wild type start site and antisense transcription from some normally unidirectional promoters. In principle, these observations are consistent with SPARC regulating either RNA synthesis or degradation. However, the balance of evidence suggests SPARC directly regulates transcription. First, our ChIP-seq datasets showed that SPARC subunits associate with RNAPII promoters in a narrow region of enrichment that matches that of RPB1. These similar enrichment patterns suggest a possible direct link between SPARC and the RNAPII transcription machinery of *T. brucei*. Such a link is supported by the interactions of CRD1, PHD6 and PWWP1 with RNAPII subunits in our proteomics analyses (Figure 1). By contrast, we did not observe any interactions between SPARC subunits and RNA decay factors. Second, SET27 loss disrupted RPB1 association with TSRs, suggesting SPARC directly or indirectly regulates transcription initiation. Taken together, these results support the conclusion that SPARC functions at *T. brucei* promoters by interacting with RNAPII to accurately demarcate transcription start sites and, in some instances, ensure unidirectional transcription (Figure 7).

**Figure 7.**
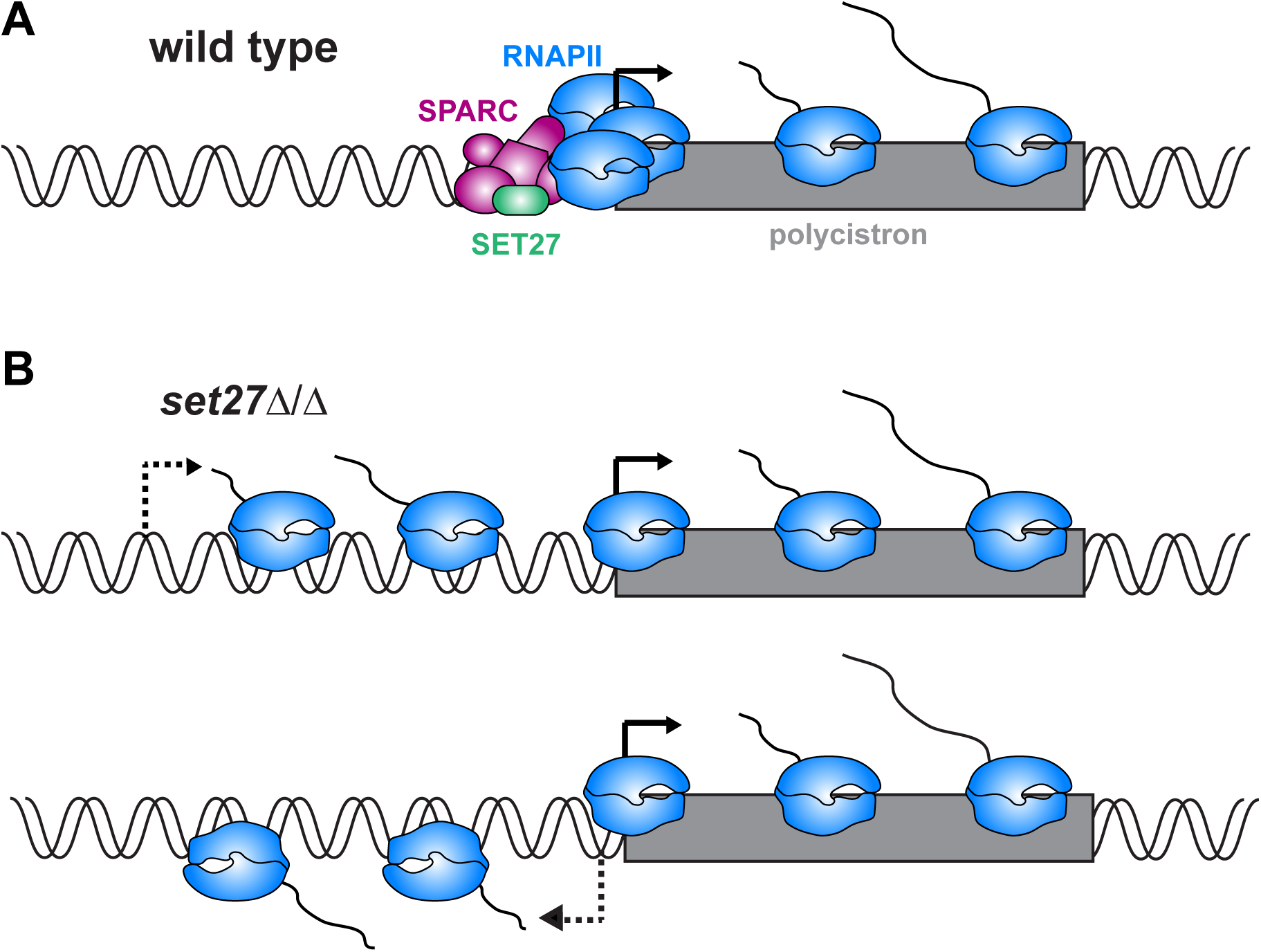
Model for SPARC-mediated definition of RNAPII transcription start sites. **(A)** In wild type cells, SPARC associates with genomic sites just upstream of polycistronic transcription units. SPARC sites coincide with promoters and regions of RNAPII enrichment. In the absence of SET27, the SPARC complex dissociates from promoters, and RNAPII enrichment is decreased. Transcription initiates upstream of the natural site (*top*), and sometimes in the reverse direction (*bottom*), effectively converting some unidirectional promoters into bidirectional promoters.

While SPARC contributes to the accuracy of transcription initiation across essentially all promoters, the abundance of most mRNAs was unaffected following SET27 loss. This probably reflects the dominant role of post-transcriptional regulation in *T. brucei* gene expression (Clayton, 2019). Such post-transcriptional buffering might have evolved primarily to regulate expressed mRNAs, leaving regions close to promoters, which are not normally transcribed, free from this regulation. This may explain why increased transcript levels in *set27*Δ/Δ were restricted to the vicinity of TSRs. However, widespread transcriptional dysregulation of mRNAs and its post-transcriptional correction might be energetically costly, potentially leading to the growth defect observed in *SET27* null cells.

Of the genes that changed expression after SET27 deletion, almost half belong to the VSG family of surface coat proteins. Bloodstream form trypanosomes normally express only a single VSG gene, whereas we detected upregulation of close to 50 VSG transcripts in *set27*Δ/Δ. The interactions of FLAG-CRD1 with VSGs in the affinity selections from *set27*Δ/Δ cells indicate that some of the normally silent VSG genes are also translated into proteins in the absence of SET27. Most of the upregulated VSG transcripts originated from subtelomeric regions, far from SPARC binding sites. We conclude that, in contrast to promoter regions, transcriptional silencing within subtelomeric regions is sensitive to loss of SPARC activity over large chromosomal domains.

Additionally, in cells lacking SET27 we observed derepression of promoter-proximal transcripts derived from retrotransposons. It is possible that a more open chromatin configuration at trypanosome TSRs allows preferential insertion of transposable elements at those locations, similar to what has been observed in other eukaryotes (Feschotte, 2008; Miao et al., 2020). In wild type cells, SPARC may help maintain genome integrity by preventing the expression of these mobile genetic elements.

In summary, we have identified SPARC, a novel promoter-associated protein complex that helps define RNAPII transcription start sites, influences transcription directionality and represses transcription across subtelomeric domains. The function of the SET27 putative histone methyltransferase at *T. brucei* TSRs is reminiscent of the role that the Set1/COMPASS complex plays at Opisthokont RNAPII promoters where it writes the H3K4me mark (Shilatifard, 2012). H3K4 methylation by Set1/COMPASS is mediated through a PHD reader module and interactions with RNAPII, influencing cryptic transcript degradation and both activation and repression of RNAPII transcription (Howe et al., 2017). Despite the divergence of histones in trypanosomes, they also exhibit H3K4 methylation that is enriched over promoter-proximal regions (Kraus et al., 2020; Wright et al., 2010). It remains to be determined which histone lysine residues, if any, are methylated by SET27, which reader proteins (i.e. CRD1, PHD6 and/or PWWP1) bind the resulting modification(s) and if their binding also contributes to promoter definition.

## MATERIALS AND METHODS

### Cell culture

Bloodstream form *T. brucei* 427 parasites were grown at 37°C and 5% CO_2_ in HMI-9 medium supplemented with Penicillin-Streptomycin (Gibco) and 10% Fetal Bovine Serum (Gibco). Procyclic form *T. brucei* 427 cells were grown at 27°C in SDM-79 medium supplemented with hemin, Penicillin-Streptomycin (Gibco) and 10% Fetal Bovine Serum (Gibco).

### Structural Bioinformatics

The presence of putative structural/functional domains in the SPARC complex subunits was identified by analyzing: (1) the amino acid sequence composition of individual subunits for remote protein homology using the HHPred web server (Söding et al., 2005), offered by the MPI Bioinformatics Tool kit (https://toolkit.tuebingen.mpg.de/tools/hhpred; Zimmermann et al., 2018; Gabler et al., 2020), (2) the three-dimensional models generated by AlphaFold (Jumper et al., 2021), an Artificial Intelligence (AI) based high accuracy structure prediction method, using ColabFod notebook on Google CoLab (Mirdita et al., 2022). AlphaFold generated three-dimensional models were further analysed using the DALI server (http://ekhidna2.biocenter.helsinki.fi/dali/; Holm and Rosenstrom, 2010) a webserver that compares a query protein structure against the available protein structures in the Protein Data Bank (PDB) to strengthen the conclusions of the HHpred analyses.

### Protein tagging

Proteins were tagged N-terminally at their endogenous loci using the pPOTv4 plasmid (Dean et al., 2015) and a derivative of the plasmid made by changing the blasticidin drug resistance marker to puromycin and the YFP tag to FLAG. Tagging constructs were made by fusion PCR and consisted of the end of the 5′ UTR of each gene, a region of the pPOTv4 plasmid containing the drug resistance gene and the tag, and the beginning of the CDS of each gene. Fusion constructs were transfected into *T. brucei* by electroporation.

### Generating knock out cell lines

The *SET27* gene deletion was performed in a YFP-RPB1 cell line (WT) via homologous recombination to replace both *SET27* alleles with hygromycin and G418 drug resistance markers. The *JBP2* gene deletions was made using a CRISPR/Cas9 toolkit developed for kinetoplastids (Beneke et al., 2017). Briefly, a pJ1339 plasmid from which Cas9 is constitutively expressed (Alves et al., 2020) was integrated into the tubulin locus to generate the J1339 WT cell line. Then, guide RNAs and repair cassettes were transfected into the J1339 WT cell line to replace both *JBP2* alleles with hygromycin and G418 drug resistance markers.

### Immunolocalization

Parasites were harvested by centrifugation, washed once with PBS and fixed with 4% paraformaldehyde (final concentration) for 10 min on ice. Fixation was then stopped by adding glycine. Fixed cells were pipetted onto polylysine slides and permeabilized with 0.1% Triton X-100. Slides were blocked with 2% BSA for 45 min at 37 °C. Blocked slides were incubated for 45 min at 37 °C first with rabbit anti-GFP primary antibody (A-11122; Thermo Fisher Scientific) diluted 1:500 in 2% BSA, and then with Alexa fluor 568 goat anti-rabbit secondary antibody (A-11036; Thermo Fisher Scientific) diluted 1:1000 in 2% BSA. DNA was stained with DAPI. Slides were imaged using a Zeiss Axio Imager microscope.

### ChIP-seq

3-5 × 10^8^ cells were harvested by centrifugation and fixed with 0.8% formaldehyde (final concentration) for 20 min at room temperature. Cells were lysed and chromatin was sheared by sonication in a Bioruptor sonicator (Diagenode) for 30 cycles (each cycle: 30 s ON, 30 s OFF; high setting). Sonication was performed in the presence of 0.2% SDS, after which sonicated samples were centrifuged to pellet cell debris and SDS in the supernatants was diluted to 0.07%. An input (0.8% of total sample) was taken before incubating the rest of the supernatant overnight with Protein G Dynabeads and 10 µg anti-GFP antibody (A-11122; Thermo Fisher Scientific). The following day, the beads were washed first in the presence of 500 mM NaCl, and then in the presence of 250 mM LiCl. The DNA was eluted from the beads using a buffer containing 50 mM Tris-HCl pH 8, 10 mM EDTA and 1% SDS. The eluted sample was treated with RNase and Proteinase K, and DNA was purified using a QIAquick PCR Purification Kit (Qiagen). ChIP-seq libraries were prepared using NEXTflex barcoded adapters (Bioo Scientific). The libraries were sequenced by the Edinburgh Clinical Research Facility on Illumina NextSeq 550 or Illumina NextSeq 2000 instruments. Subsequent ChIP-seq data analysis was based on three replicates for CRD1 (BF), JBP2 (BF), and SET27 (PF), and two replicates for the other YFP-tagged proteins and for the untagged control in BF and PF cells. ChIP-seq data for CRD1 (BF), SET27 (BF) and the untagged control (BF) was taken from Staneva et al., 2021.

### ChIP-seq data analysis

ChIP-seq reads were first de-duplicated with pyFastqDuplicateRemover (Webb et al., 2018) and then aligned to the Tb427v9.2 genome (Muller et al., 2018) with Bowtie 2 (Langmead and Salzberg, 2012). Each ChIP sample was normalized to its input (as a ratio) and to library size (as counts per million). ChIP-seq data was visualized using IGV (Robinson et al., 2011). Metagene plots and heatmaps were generated as follows. First, CRD1 peaks were called using a combination of MACS2 (Feng et al., 2012) and manual filtering of false positives including peaks absent in some CRD1 replicates, peaks called in the untagged control cell line, and peaks with IP/input ratios < 6.5. Then, 20 kb regions centered around each CRD1 summit were divided into 50 bp windows. Metagene plots were generated by adding the normalized reads in each 50 bp window and then representing them as a density around CRD1. The heatmaps show normalized reads around individual CRD1 peaks.

### RNA-seq

1-5 × 10^7^ parasites were harvested by centrifugation and RNA was extracted from cells using Trizol (Thermo Fisher Scientific) according to the manufacturer’s protocol. RNA samples were treated with DNase followed by purification with Phenol/Chloroform/Isoamyl alcohol pH 4. Libraries were prepared and sequenced by the Edinburgh Clinical Research Facility using NEBNEXT Ultra II Directional RNA Library Prep kit (NEB) and Poly-A mRNA magnetic isolation module (NEB). The libraries were sequenced on Illumina NextSeq 550 or Illumina NextSeq 2000 instruments. Subsequent RNA-seq data analysis was based on two replicates for the wild type, *jbp2*Δ/+ and *jbp2*Δ/Δ cell lines, and three replicates for the *set27*Δ/+ and *set27*Δ/Δ cell lines

### RNA-seq data analysis

RNA-seq reads were aligned to the Tb427v9.2 genome (Muller et al., 2018) with Bowtie 2 (Langmead and Salzberg, 2012). Aligned reads were separated into those originating from the plus or the minus DNA strand, normalized to library size (as counts per million) and visualized using IGV (Robinson et al., 2011). The differential expression analysis was performed as follows. First, raw non-normalized reads overlapping mRNAs annotated in the Tb427v9.2 genome were counted using featureCounts (Liao et al., 2014). Reads overlapping several mRNAs were assigned to the mRNA with the largest number of overlapping bases. Differential expression analysis was then performed using edgeR (Robinson et al., 2010) utilizing the TMM normalization method (Robinson and Oshlack, 2010). The filterByExpr (min.count=4; min.total.count=20) function was applied to filter out genes with insufficient counts for performing the statistical analysis.

### Affinity selections and mass spectrometry

4 × 10^8^ cells were harvested by centrifugation and lysed in a buffer containing 50 mM Tris pH 8, 150 mM KCl and 0.2% NP-40. Lysates were sonicated in a Bioruptor (Diagenode) sonicator for 3 cycles (each cycle: 12 s ON, 12 s OFF; high setting). Sonicated samples were centrifuged and the supernatant (soluble fraction) was incubated for 1 h at 4°C with beads crosslinked to anti-GFP antibody (11814460001; Roche) or to M2 anti-FLAG antibody (F1804; Sigma-Aldrich). The beads were then washed three times with lysis buffer. Proteins were eluted from the beads using RapiGest surfactant (Waters) at 55°C for 15 min. Proteins were digested by filter-aided sample preparation (FASP; Wiśniewski et al., 2009). Briefly, protein samples were reduced using DTT, then denatured with 8 M Urea in Vivakon spin column cartridges. Samples were treated with 0.05 M iodoacetamide and digested overnight with 0.5 μg MS Grade Pierce Trypsin Protease (Thermo Fisher Scientific). Digested samples were desalted using stage tips (Rappsilber et al., 2007) and resuspended in 0.1% trifluoroacetic acid for mass spectrometry analysis. LC-MS/MS was performed as described previously (Staneva et al., 2021). The results were processed using the Perseus software (Tyanova et al., 2016) and are based on three biological replicates per sample. The interaction network of the analyzed proteins was created using Cytoscape (Shannon et al., 2003).

## DATA ACCESS

The sequencing data generated in this study can be accessed on the NCBI Gene Expression Omnibus (GEO; https://www.ncbi.nlm.nih.gov/geo/) with accession number GSE208037.

## ACKNOWLEDGEMENTS

We thank Julie Young for supplying the HMI-9 media used during this project. We would also like to thank Richard Clark, Angie Fawkes, Audrey Coutts and Tamara Gilchrist from the Edinburgh Wellcome Clinical Research Facility for sequencing services as well as Shaun Webb from the WCB bioinformatics core facility for maintaining the servers we used for processing sequencing data. This work was funded by an MRC Research Grant awarded to R.C.A and K.R.M and supporting D.P.S (MR/T04702X/1), a Wellcome Investigator Award to K.R.M. (22171), a Wellcome Principal Research Fellowship to R.C.A. supporting D.P.S. and T.A (200885; 224358), a Wellcome Principal Research Fellowship to D.T. supporting S.B. (222516), a Wellcome Senior Research Fellowship to A.A.J. (202811), a Wellcome Instrument grant to J.R. (108504), and core funding for the Wellcome Centre for Cell Biology (203149) supporting C.S.

---

**Supplemental Figure S1.**
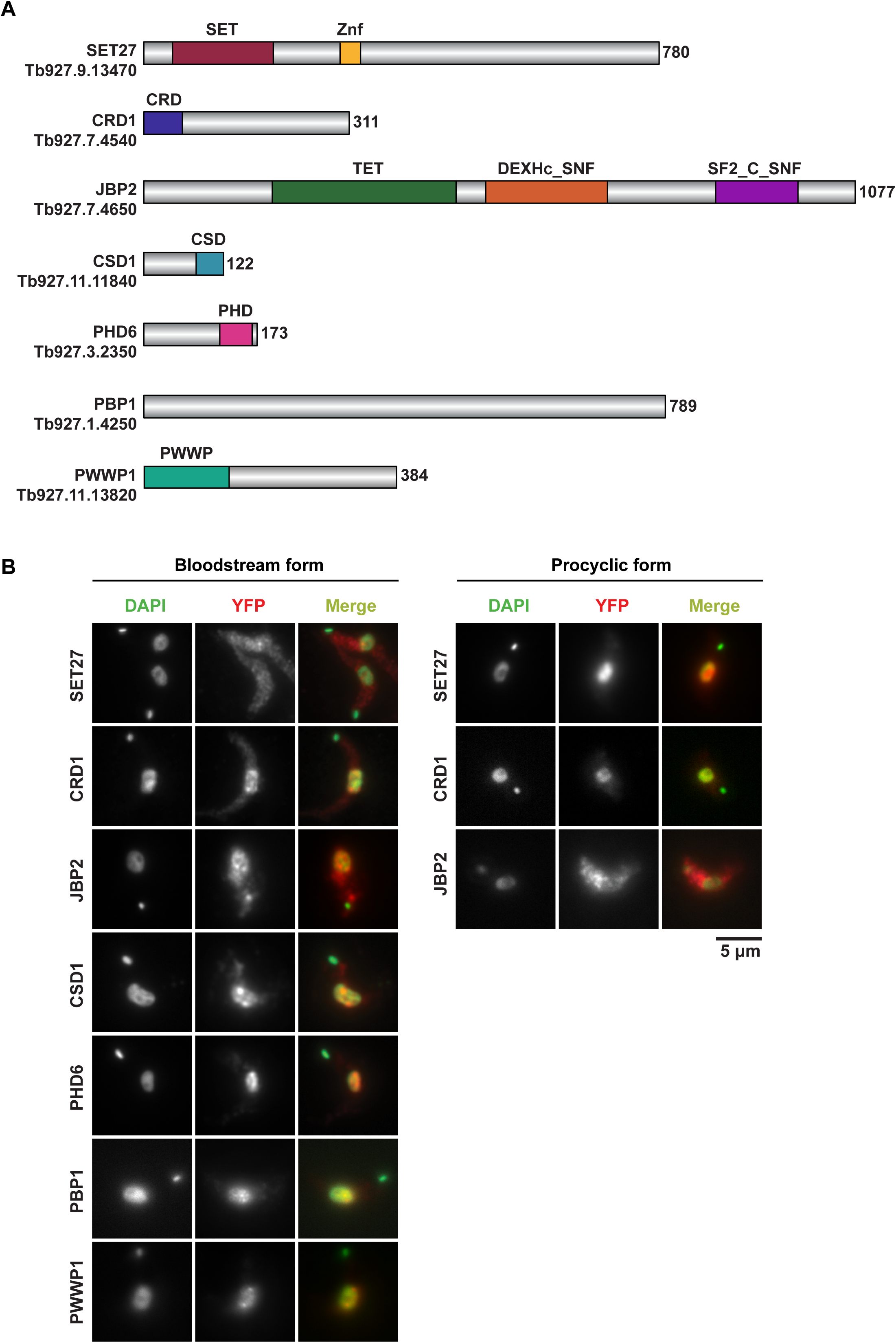
Domain architecture and localization of SPARC complex subunits. Related to Figure 1. **(A)** Conserved domains and sequences of core SPARC complex subunits and JBP2. **(B)** Immunolocalization of YFP-tagged core SPARC components and JBP2 in bloodstream and procyclic form cells. In merge panels, YFP-tagged proteins are shown in red and DAPI-stained DNA is in green.

**Supplemental Figure S2.**
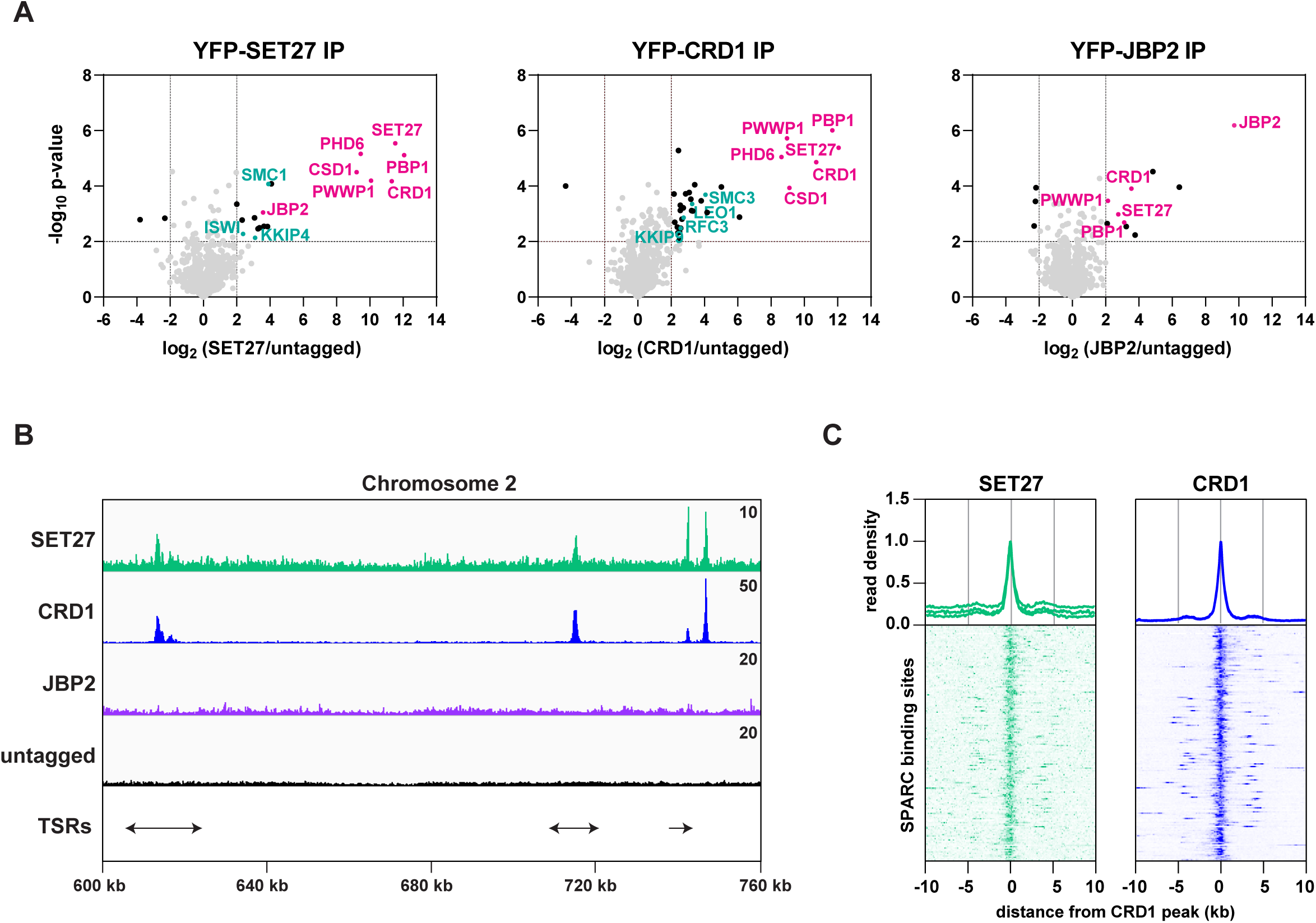
SPARC is present in procyclic form cells. Related to Figure 1 and Figure 2. **(A)** SET27, CRD1 and JBP2 were YPF-tagged in PF cells, and analyzed by affinity selection and LC-MS/MS to identify protein interaction partners. Volcano plots are based on 3 biological replicates. Cutoffs used for significance: log2 fold change (tagged/untagged) >2 or <-2 and Student’s t-test p-value < 0.01. See Supplemental Table S1 for a complete list of proteins enriched in each affinity selection. Core SPARC components and the auxiliary subunit JBP2 are shown in pink, and other proteins of interest are shown in teal. **(B)** A region of Chromosome 2 (same as in Figure 2A) is shown with ChIP-seq reads mapped for the indicated proteins. A single replicate is shown for each protein. ChIP-seq performed in cells lacking any tagged protein (untagged) was included as a negative control. Tracks are scaled separately as fragments per million (the exact value is indicated in the top-right corner of each track). The position of single and double transcription start regions (sTSRs and dTSRs) are shown below with arrows indicating the direction of transcription. **(C)** Enrichment profiles for SPARC components. CRD1 is used as a reference because it has the most prominent peaks at TSRs. The metagene plots (*top*) show normalized read density around all CRD1 peak summits, with each replicate plotted separately. The heatmaps (*bottom*) show protein density around 177 individual CRD1 peaks. Each heatmap shows the average of at least 2 replicates.

**Supplemental Figure S3.**
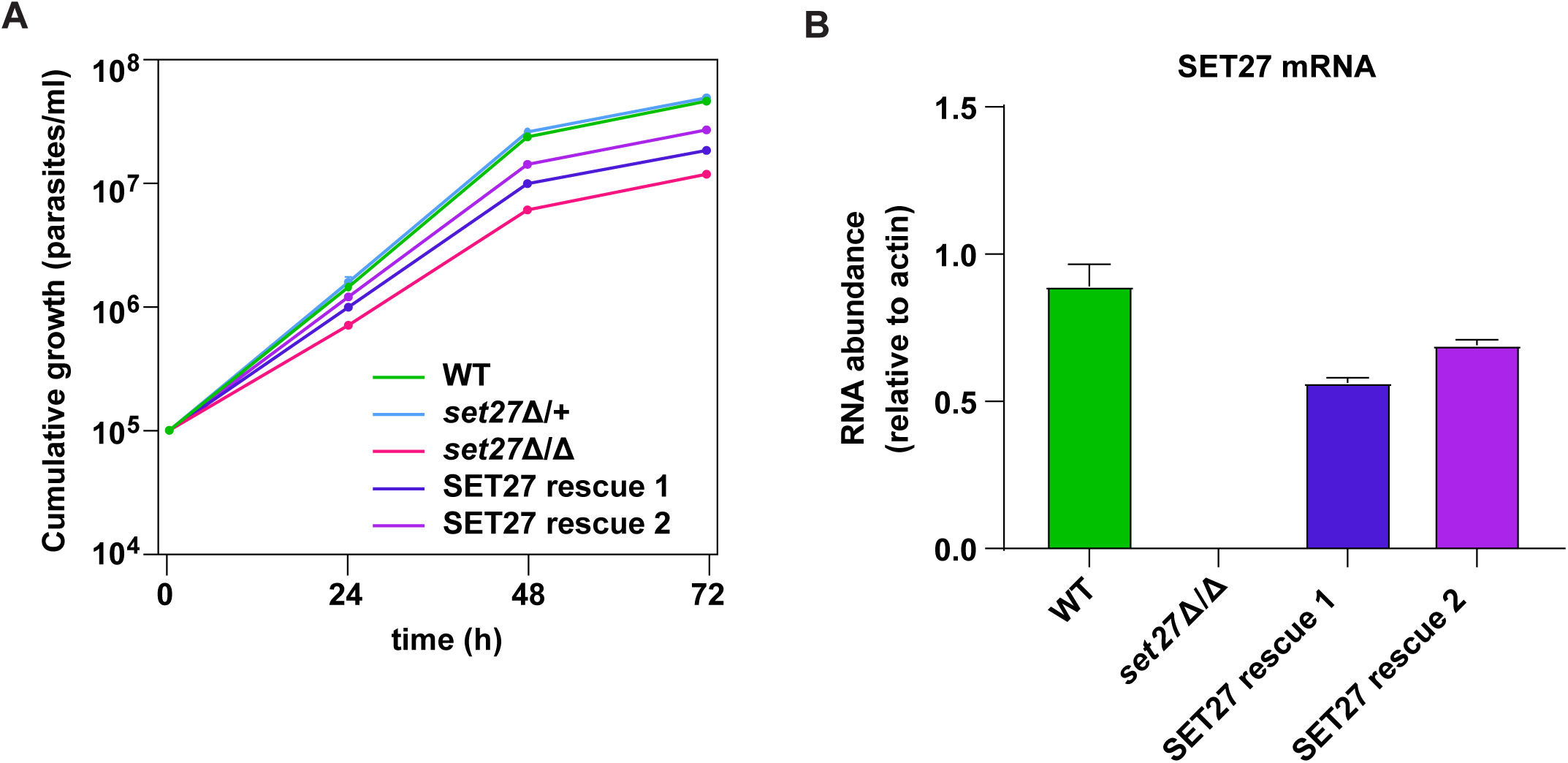
Validation of the SET27 knockout. Related to Figure 3. **(A)** Growth curves for wild type, *set27*Δ/+, *set27*Δ/Δ, and the two *SET27* rescue clones. Error bars: Standard Deviation (SD) of 3 biological replicates. Cells lacking the *SET27* gene (*set27*Δ/Δ) grew substantially slower than WT or *set27*Δ/+ cells (t test p-value < 0.0001). SET27 rescue clones 1 and 2 also grew significantly slower than WT or *set27*Δ/+ cells (t test p-value < 0.01) but faster than *set27*Δ/Δ cells (t test p-value < 0.01). Average doubling times: 6.2 h (wild type), 6.0 h (*set27*Δ/+), 8.3 h (*set27*Δ/Δ), 7.4 h (*SET27* rescue clone 1) and 6.8 h (*SET27* rescue clone 2). **(B)** SET27 mRNA levels assessed by RT-qPCR in wild type, set27Δ/Δ and the two *SET27* rescue clones. Error bars: Standard Deviation (SD) of 2 biological replicates.

**Supplemental Figure S4.**
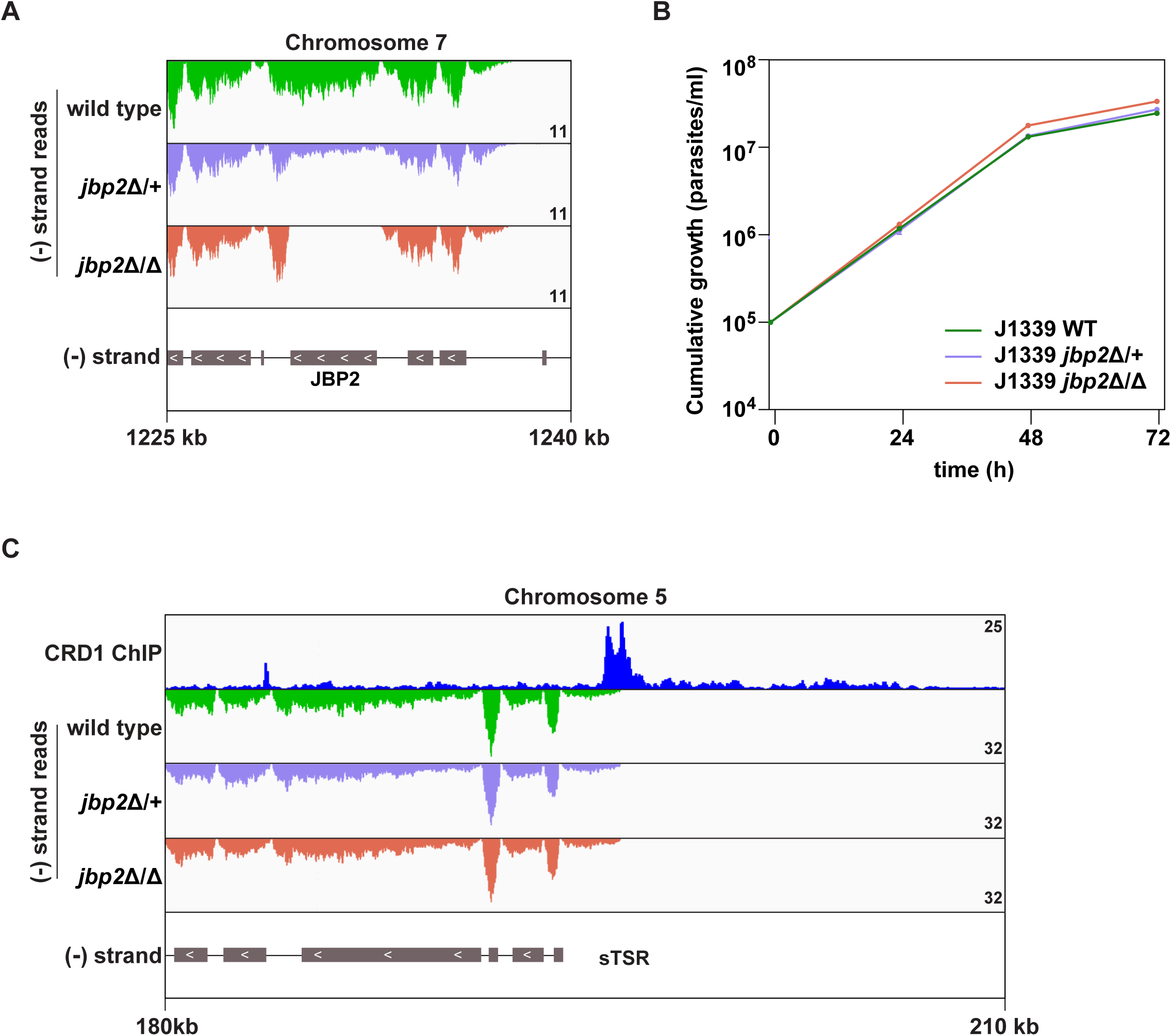
Validation of the JBP2 knockout. Related to Figure 3. **(A)** Tracks showing the distribution of RNA-seq reads in the vicinity of the *JBP2* gene in wild type, *jbp2*Δ/+ and *jbp2*Δ/Δ cells. All tracks are scaled identically (number of reads shown in the bottom right corner of each track). **(B)** Growth curves for J1339 wild type, *jbp2*Δ/+ and *jbp2*Δ/Δ cell lines. Error bars: Standard Deviation (SD) of 3 biological replicates. Cells lacking the *JBP2* gene (*jbp2*Δ/Δ) grew slightly faster than WT cells (t test p-value < 0.05) and about the same as *jbp2*Δ/+ cells (t test p-value = 0.05). Average doubling times: 6.9 h (J1339 WT), 6.8 h (J1339 *jbp2*Δ/+) and 6.5 h (J1339 *jbp2*Δ/Δ). Note that, in contrast to SET27 knockouts, JBP2 deletions were made via CRISPR/Cas9 in a cell line with integrated pJ1339 plasmid (see Materials and Methods). **(C)** Tracks showing the distribution of RNA-seq reads in the presence (wild type and *jbp2*Δ/+) or absence of JBP2 (*jbp2*Δ/Δ) around the same sTSR shown in Figure 4A (*left*). CRD1 ChIP (*top track*) is included to mark the position of SPARC sites. ORFs are indicated by grey boxes, and directionality is shown with inset white arrows. Genes present within a single polycistron are connected with a thin black line.

